# TRIP13 fosters both transcriptional silencing and DSB repair during meiosis

**DOI:** 10.1101/2025.10.20.683374

**Authors:** Marina Marcet-Ortega, Ana Martínez-Marchal, Andros Maldonado-Linares, Mercè Encinas, Anna Esteve Codina, Maria Jasin, Scott Keeney, Ignasi Roig

## Abstract

During meiosis, the coordinated suppression of global gene transcription and induction of double-strand breaks (DSBs) genome wide is crucial for successful gamete formation. The AAA+ ATPase TRIP13 is a master meiotic regulator involved in numerous cellular processes, including DSB repair, organization of the meiotic chromosome axis, and meiotic silencing of unsynapsed axes. Here, we examine the involvement of TRIP13 in transcription suppression and DSB repair during meiotic prophase I in mice. Because TRIP13 is required for silencing of sex chromosomes at pachynema, we hypothesized that it may also be involved in the global transcription suppression that begins at the onset of meiotic prophase. Indeed, as predicted, we observed increased incorporation of 5-ethynyl uridine in early stages of meiotic prophase I and a higher presence of the active form of RNA polymerase II in *Trip13* mutant spermatocytes. RNA sequencing data revealed that more than two-thirds of upregulated transcripts came from the sex chromosomes. Given this more general role in transcription silencing, we also tested whether TRIP13 promotes the transcription block that causes meiotic arrest in synapsis-deficient oocytes. However, *Trip13* ablation did not rescue *Spo11^-/-^* oocytes, which experience asynapsis without DSB repair defects. Instead, *Trip13* mutation did increase the number of oocytes in *Dmc1^-/-^ Chk2^-/-^* mice, in which DSBs persist unrepaired. This rescue seems attributable to a previously unknown function of TRIP13 in preventing DSB repair by non-canonical pathways, as we observed fewer homologous recombination markers along with an increased presence of non-homologous end joining markers in the triple mutant cells. These findings underscore TRIP13’s multifaceted role in transcription and DSB repair regulation during meiosis, highlighting its importance and potential for further meiotic regulation and fertility research.

## Introduction

Meiotic prophase I is a highly orchestrated stage during gametogenesis, marked by extensive chromatin remodeling, homologous chromosome pairing, synapsis, and recombination. In mammals, these processes are initiated by the expression of meiosis-inducing factors such as STRA8 and MEIOSIN, which are activated by retinoic acid signaling (Anderson et al., 2008; Ishiguro et al., 2020). These transcriptional regulators promote the expression of key meiotic genes such as *Spo11* (Kojima et al., 2019), which encodes the topoisomerase-like enzyme responsible for generating programmed DNA double-stranded beaks (DSBs) (Keeney et al., 1999; Baudat et al., 2000).

Repair of these DSBs by homologous recombination facilitates the pairing of homologous chromosomes, a process stabilized by the assembly of the synaptonemal complex (SC), a tripartite protein structure that bridges the chromosome axes and provides a scaffold for recombination (Bolcun-Filas and Handel, 2018; Huang and Roig, 2023). Meiotic prophase I is traditionally subdivided according to SC formation. At leptonema, each homolog organizes around a chromosome core, also known as an axial element. During zygonema, homologs pair and synapse. At pachynema, a fully formed SC links the entire length of the homologous chromosomes. At diplonema, the SC disassembles, but the homologs remain linked by chiasmata, which are the cytological manifestation of crossovers.

In parallel with the induction of DSBs by SPO11, and as a consequence of the initiation of the meiotic program mediated by STRA8 (Bellutti et al., 2024), transcription is globally suppressed at the onset of prophase I (Monesi, 1964; Page et al., 2012). Transcription activity resumes at mid-pachynema, once synapsis is complete and most DSBs are repaired. Timely and accurate completion of homologous recombination is essential for progression through pachynema and for the maintenance of genomic integrity (Bolcun-Filas et al., 2014; Pacheco et al., 2015; Marcet-Ortega et al., 2017; Martínez-Marchal et al., 2020).

In pachytene spermatocytes, all homologous pairs are fully synapsed except for the sex chromosomes, which share only a small region of homology called the pseudoautosomal region. Thus, most of the X and Y chromosomes remain unsynapsed for most of pachynema. While autosomes resume transcriptional activity at this stage, the unsynapsed regions of the sex chromosomes remain transcriptionally silenced through heterochromatinization, a process known as meiotic sex chromosome inactivation (MSCI)(Turner et al., 2004). MSCI is considered a specialized manifestation of the broader mechanism of meiotic silencing of unsynapsed chromosomes (MSUC), which is initiated during zygonema (Burgoyne et al., 2009). Proper silencing of the sex chromosomes is essential for meiotic progression and male fertility (Royo et al., 2010).

We previously demonstrated that the AAA+ ATPase TRIP13 is a key regulator of meiotic prophase, orchestrating multiple aspects of chromosome dynamics in spermatocytes (Roig et al., 2010), including the establishment of MSCI (Pacheco et al., 2015; Marcet-Ortega et al., 2017). Given the proposed link between MSUC and global transcriptional repression that occurs at meiotic entry (Page et al., 2012), we hypothesized that TRIP13 may also contribute to transcriptional regulation at the onset of meiosis. To investigate this, we assessed transcriptional activity in early prophase I in both wild-type and *Trip13* mutant spermatocytes. Additionally, we examined whether TRIP13 is required to promote meiotic arrest in synapsis-deficient oocytes, where transcriptional dysregulation is known to occur.

Our findings indicate that TRIP13 is essential for sex chromosome silencing in spermatocytes but is dispensable for oocyte arrest in response to asynapsis. Moreover, we provide evidence that TRIP13 plays a previously unrecognized role in preventing DSB repair via non-canonical pathways during meiosis, further highlighting its multifaceted function in maintaining genome integrity during gametogenesis.

## Results

### TRIP13 regulates gene expression of the sex chromosomes and autosomes during meiotic prophase I

To test for a role for TRIP13 in global meiotic gene expression, we cytologically detected newly synthesized RNA using 5-ethynyl uridine (EU) incorporation in spermatocyte cell suspensions followed by fluorophore labeling by click chemistry under conditions that preserved nuclear structure and cell volume (Fig. 1A). To classify meiotic prophase stages, cells were also immunofluorescently stained to detect SYCP3, a component of the lateral element of the SC, and γH2AX, a marker for DNA damage.

**Figure 1.**
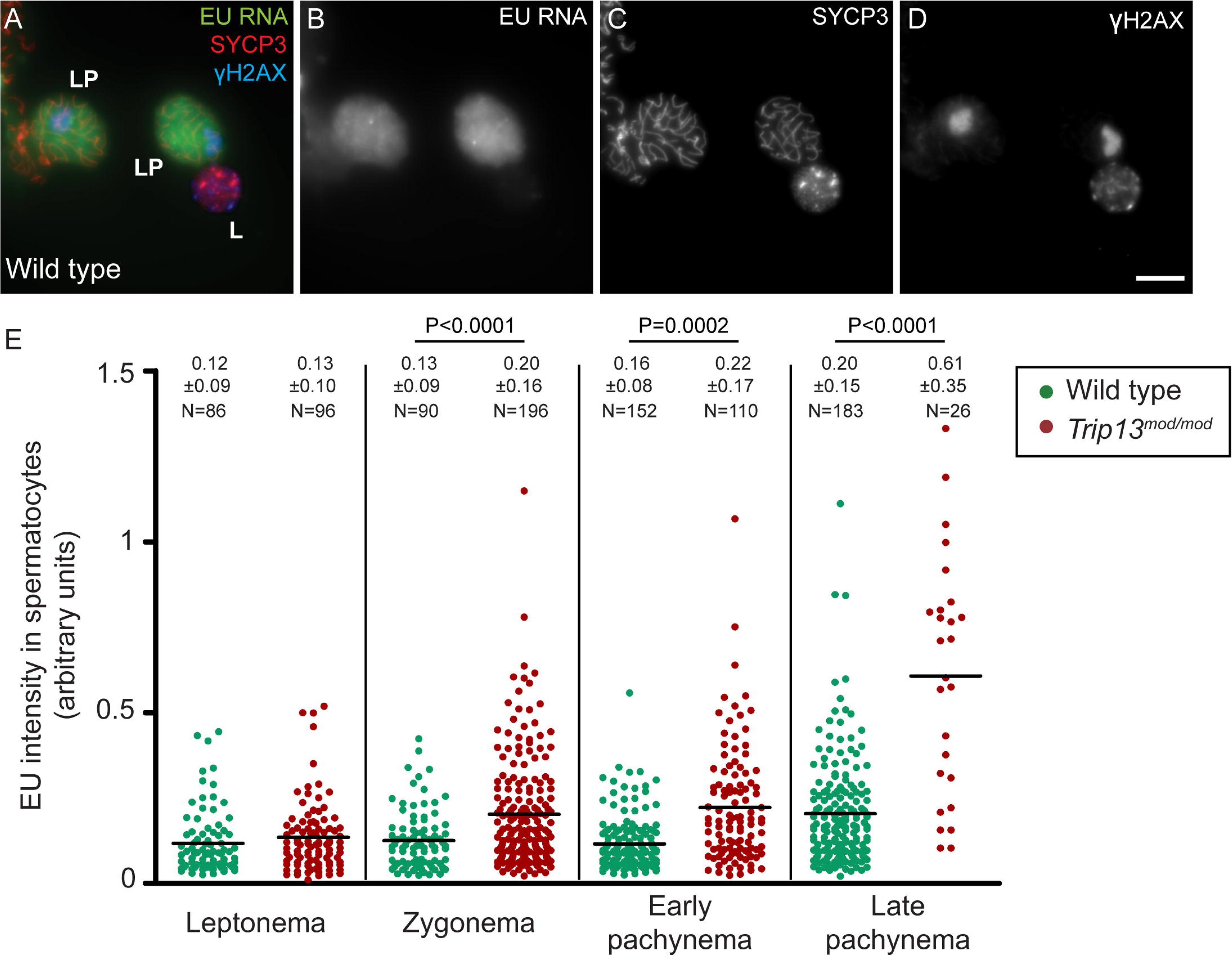
TRIP13 is required to prevent transcription in the meiotic prophase. (A) Representative image (sum projection of stack images) of EU-treated wild-type late pachytene (LP) and leptotene (L) spermatocytes. The scale bar represents 10 μm. (B) Quantification of the EU intensity in *Trip13^mod/mod^* and wild-type spermatocytes at the indicated prophase stages. Horizontal lines represent means. Mean ± SD and the number of cells counted (N) are indicated above the graph. P values are from t tests. A total of five mice were analyzed per each genotype.

This approach confirmed that gene expression in wild type is low in early meiotic prophase I and increases at mid-pachynema, as previously described (Fig. 1A,B and Monesi, 1964; Page et al., 2012). In contrast, mice homozygous for a hypomorphic *Trip13* mutation (*Trip13^mod/mod^*, for “moderate”) that significantly reduces TRIP13 expression (Roig et al., 2010), presented significantly higher EU signal compared to wild-type cells at zygonema and pachynema (Fig. 1B). We observed a similar effect in spermatocytes from another *Trip13* mutant line that presents a more penetrant phenotype (*Trip13^sev/sev^*, for “severe”; Fig. S1).

The increased transcription levels appears not to be caused by the arrest triggered by defective recombination that occurs in *Trip13* mutants because *Dmc1^-/-^*spermatocytes, which cannot repair DSBs and arrest at a similar stage (Pittman et al., 1998), did not show similarly increased EU incorporation (Fig. S1). The arrest triggered in *Spo11^-/-^* spermatocytes (Baudat et al., 2000) also did not result in increased EU labelling (Fig. S1).

To confirm these findings, we examined the presence of the active form of RNA polymerase II, detected by immunostaining with an antibody specific for phosphorylation on Ser-2 within the YSPTSPS repeat in the RNA polymerase II C-terminal domain (Han et al., 2016) (pRNAP). pRNAP staining in wild-type spermatocytes showed diffuse nuclear signal plus a few small clumps (Fig. 2A). By contrast, *Trip13* mutants formed more and larger clumps, particularly evident in zygonema (Fig. 2B,C). Furthermore, there was a significant increase in the pRNAP staining intensity in *Trip13^mod/mod^* and *Trip13^sev/sev^* mutants at zygonema (Fig. 2D,E).

**Figure 2.**
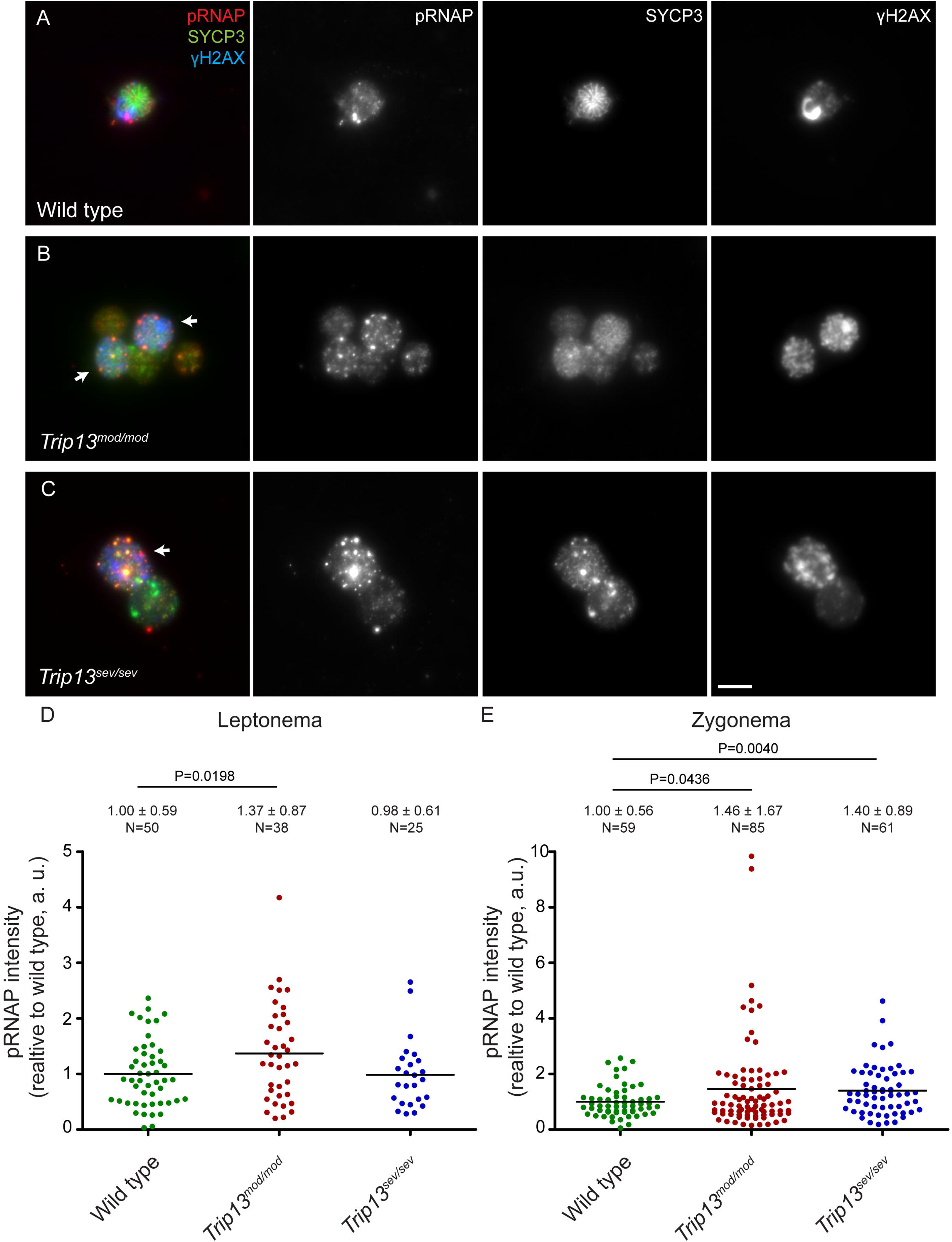
TRIP13 inhibits activation of RNA polymerase II during early meiotic prophase. **I.** (A–C) pRNAP levels in representative zygotene spermatocytes (arrows) from wild type (A), *Trip13^mod/mod^*(B) and *Trip13^sev/sev^* (C). The sum projection of stack images is shown. The scale bar in C represents 10 μm and applies to all panels. (D, E) Quantification of pRNAP intensity (arbitrary units, a.u.) in leptotene (D) and zygotene (E) spermatocytes of the indicated genotypes, relative to the average in wild type. Horizontal lines represent means. Means ± SD and the number of counted cells (N) are indicated above the graph. P values are from t tests. Four, three and one mice were analyzed for wild type, *Trip13^mod/mod^*and *Trip13^sev/sev^*, respectively.

Taken together, these results suggest that transcriptional activity is increased in early prophase I in *Trip13* mutants. To better characterize this increase, we performed RNA sequencing (RNAseq) on testes of juvenile wild-type and *Trip13^mod/mod^* mice. Overall, we found 683 misregulated genes in *Trip13* mutant samples, 462 downregulated and 221 upregulated (Fig. 3A). We validated these results by quantitative real-time PCR analysis for three overexpressed genes (*Xaf1*, *Tktl1,* and *Zfy2*, Fig. S2A) and three downregulated genes (*Atp8b3*, *Qrich2,* and *Trip13*, Fig. S2B). Gene ontology analysis of the misregulated genes revealed the overrepresentation of several spermatogenesis-related processes, such as axoneme assembly, cilium-related biological processes, spermatogenesis, or sexual reproduction (Fig. 3B).

**Figure 3.**
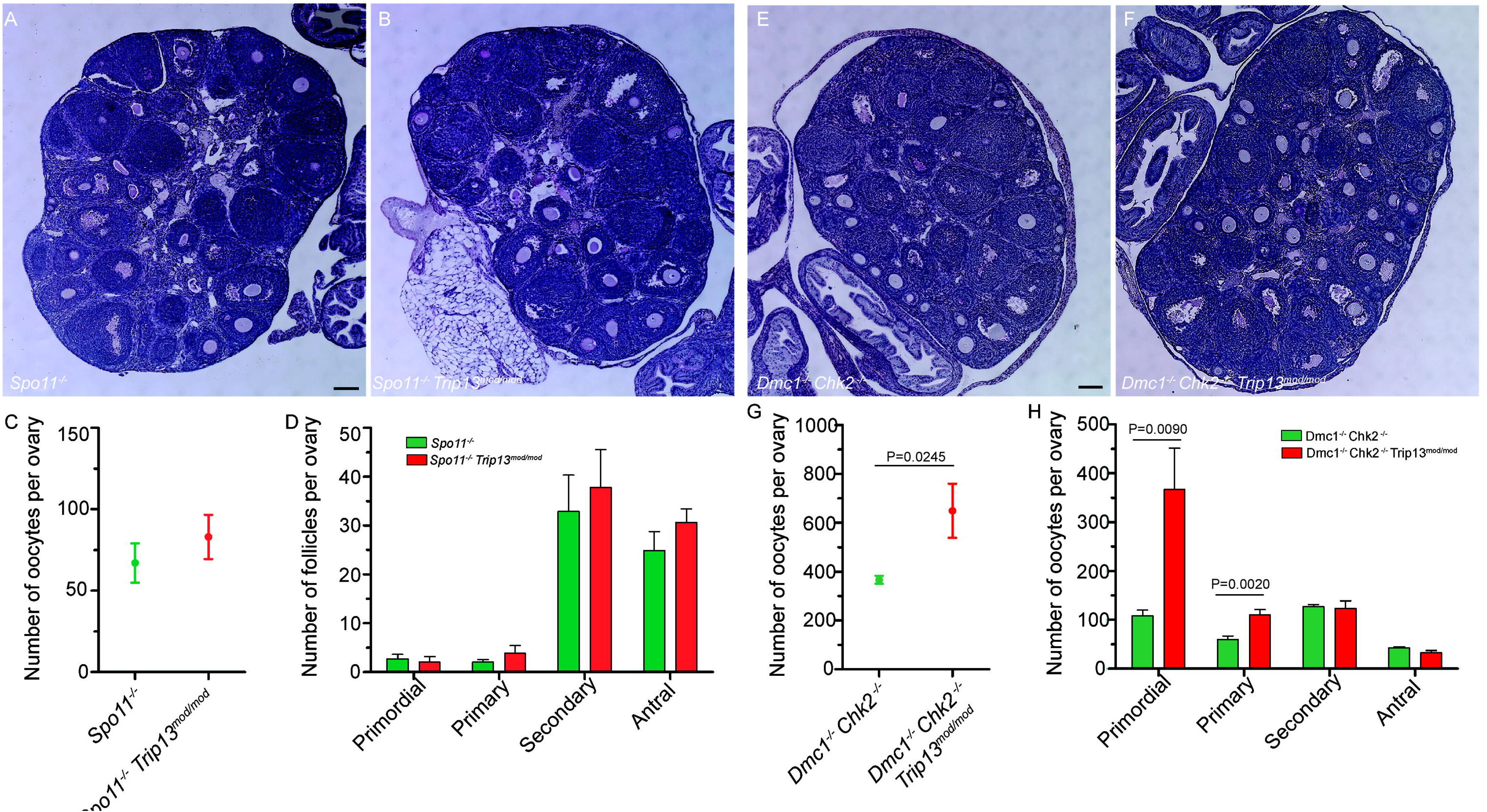
TRIP13 regulates global gene expression during meiosis. (A) Volcano plot showing all differentially expressed genes (DEGs) (FDR<0.01; red) in 14 dpp wild type testis compared to 14 dpp *Trip13^mod/mod^* testis (N=3 wild type and 3 *Trip13^mod/mod^* mice). The names of the top ten DEGs are shown. Genes with a positive fold change represent genes at higher level in the wild type. Genes with a negative fold change represent genes expressed at higher level in the *Trip13^mod/mod^* testis. (B) Gene Ontology analysis of the DEGs. The top 15 misregulated biological processes according to their fold enrichment are displayed. The number of genes in each process and their FDR are also shown. (C) Representation of the location along the genome of all upregulated and downregulated genes in *Trip13^mod/mod^* testis.

Interestingly, while genes that were downregulated in the mutant are dispersed across all autosomes in a frequency that corresponded well with gene content (Table S1), almost 70% of the upregulated genes are located on the sex chromosomes (Fig. 3C), which is 15 and 5 times more than expected based on the X and Y chromosome gene content, respectively. The more pronounced effect of TRIP13 deficiency on transcription of the sex chromosomes confirms its previously described role in MSCI (Pacheco et al., 2015; Marcet-Ortega et al., 2017).

### *Trip13* mutation rescues the oocyte number in *Dmc1^-/-^ Chk2^-/-^* mice

Since MSCI is thought to be related to MSUC, and because MSUC eliminates most oocytes of mice with defective synapsis (Cloutier et al., 2015; Martínez-Marchal et al., 2020), we hypothesized that TRIP13 deficiency might rescue the oocyte-loss phenotype in synapsis-defective mice. To test this, we analyzed the number of oocytes present in 30 dpp females of two different genotypes that have synapsis defects: *Spo11^-/-^* mice, which cannot form DSBs (Baudat et al., 2000; Romanienko and Camerini-Otero, 2000), and *Dmc1^-/-^ Chk2^-/-^* mice, which lack the meiotic recombinase DMC1 and cannot perform the homology search necessary for homologous synapsis (Pittman et al., 1998; Yoshida et al., 1998). Oocytes of *Dmc1^-/-^ Chk2^-/-^* mice accumulate recombination intermediates and fail to complete synapsis but do not arrest because they lack the DNA damage response kinase CHK2 (Bolcun-Filas et al., 2014; Rinaldi et al., 2017).

Both synapsis-defective mutants presented significantly fewer oocytes than wild-type mice (Table S2, Fig. 4 and Martínez-Marchal et al., 2020), with *Spo11^-/-^* having fewer oocytes than *Dmc1^-/-^ Chk2^-/-^* mice because *Chk2* ablation partly rescues the oocyte population in recombination-deficient mice (Bolcun-Filas et al., 2014; Martínez-Marchal et al., 2020). Introducing the *Trip13* mutation significantly increased oocyte numbers in the *Dmc1^-/-^ Chk2^-/-^* background (P=0.0245, t test, Table S2, Fig. 4E,G,H). Notably, the *Dmc1^-/-^ Chk2^-/-^ Trip13^mod/mod^* triple mutant females had significantly more primordial follicles (P=0.009, t test, Table S2, Fig. 4H), which correspond to the stock of oocytes females will use during their reproductive life, suggesting that the rescue by *Trip13* mutation occurred before follicle formation or while oocytes were arrested at this stage.

**Figure 4.**
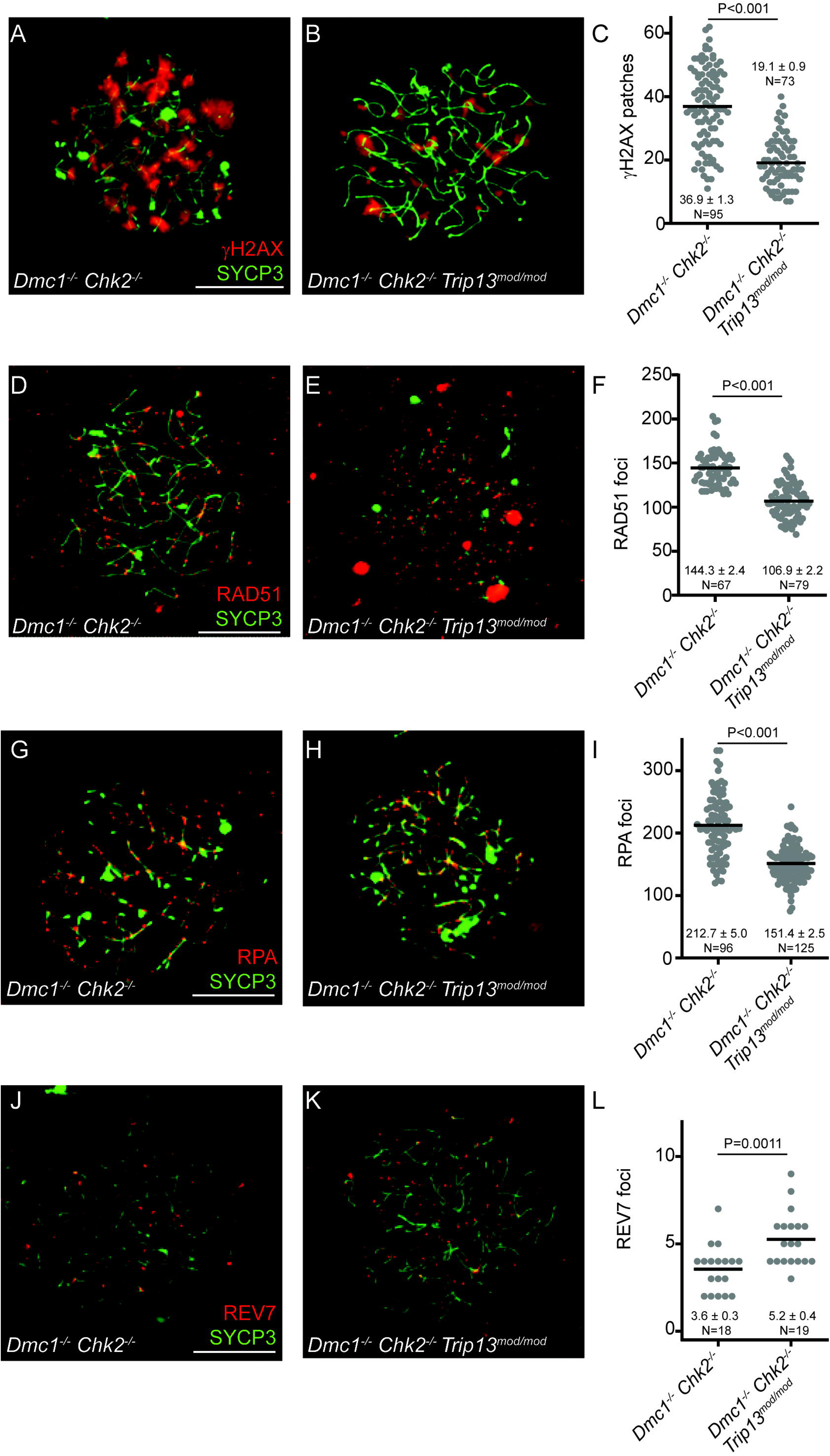
*Trip13* mutation rescues *Dmc1 Chk2* mutant oocyte arrest, but not the *Spo11* mutant one. (A,B) Histological sections of *Spo11^-/-^* (A) and *Spo11^-/-^ Trip13^mod/mod^* (B) ovaries stained with PAS-Hematoxylin. The scale bar represents 100 μm and applies to both images. (C) Number of oocytes per ovary in 30 dpp ovaries. The round symbols represent the mean and the lines, the SEM, N=8 mice per genotype. (D) Number of oocytes classified for each follicle type. The bars represent means ± SEM. (E-H) Histological sections of *Dmc1^-/-^ Chk2^-/-^* (E) and *Dmc1^-/-^ Chk2^-/-^ Trip13^mod/mod^* (F) ovaries plus quantification of total oocytes (G) and follicle types (H), presented as described for panels A–D. N=8 mice per genotype. The P values are from t tests.

In contrast, *Trip13* mutation did not significantly affect oocyte number in *Spo11^-/-^* mice (P=0.383, t test, Table S2, Fig. 4). The fact that *Trip13* deficiency does not rescue both synapsis-deficient mutant models suggests that the rescue observed in *Dmc1^-/-^ Chk2^-/-^ Trip13^mod/mod^* female mice may be attributable to reasons other than simply ameliorating the consequences of synapsis defects.

### *Trip13* mutation promotes DSB repair in *Dmc1^-/-^ Chk2^-/-^* cells

TRIP13 has been implicated in DSB repair pathway choice favoring homologous recombination over non-homologous end joining (NHEJ) in somatic cells (Clairmont et al., 2020). TRIP13 does so by remodeling the NHEJ-promoting factor REV7, a HORMA domain-containing protein that is part of the shieldin complex, which favors NHEJ by inhibiting DSB end resection (Ghezraoui et al., 2018; Gupta et al., 2018; Mirman et al., 2018; Noordermeer et al., 2018; Clairmont et al., 2020; Sarangi et al., 2020). Thus, in the absence of TRIP13, DSBs are more readily repaired by NHEJ (Clairmont et al., 2020).

We therefore hypothesized that TRIP13 deficiency might partially rescue *Dmc1^-/-^ Chk2^-/-^* cells by facilitating DSB repair. To test this, we analyzed the presence of unrepaired DSBs by staining for γH2AX in pachytene-like spermatocytes of double (*Dmc1^-/-^ Chk2^-/-^*) and triple (*Dmc1^-/-^ Chk2^-/-^ Trip13^mod/mod^*) mutant mice. We examined spermatocytes instead of oocytes due to the difficulty of obtaining double and especially triple mutant fetal oocytes. Consistent with prediction, the triple mutant spermatocytes had, on average, about half the number of γH2AX patches per cell (Fig. 5A-C), suggesting that TRIP13 deficiency allowed more DSBs to be repaired.

**Figure 5.**
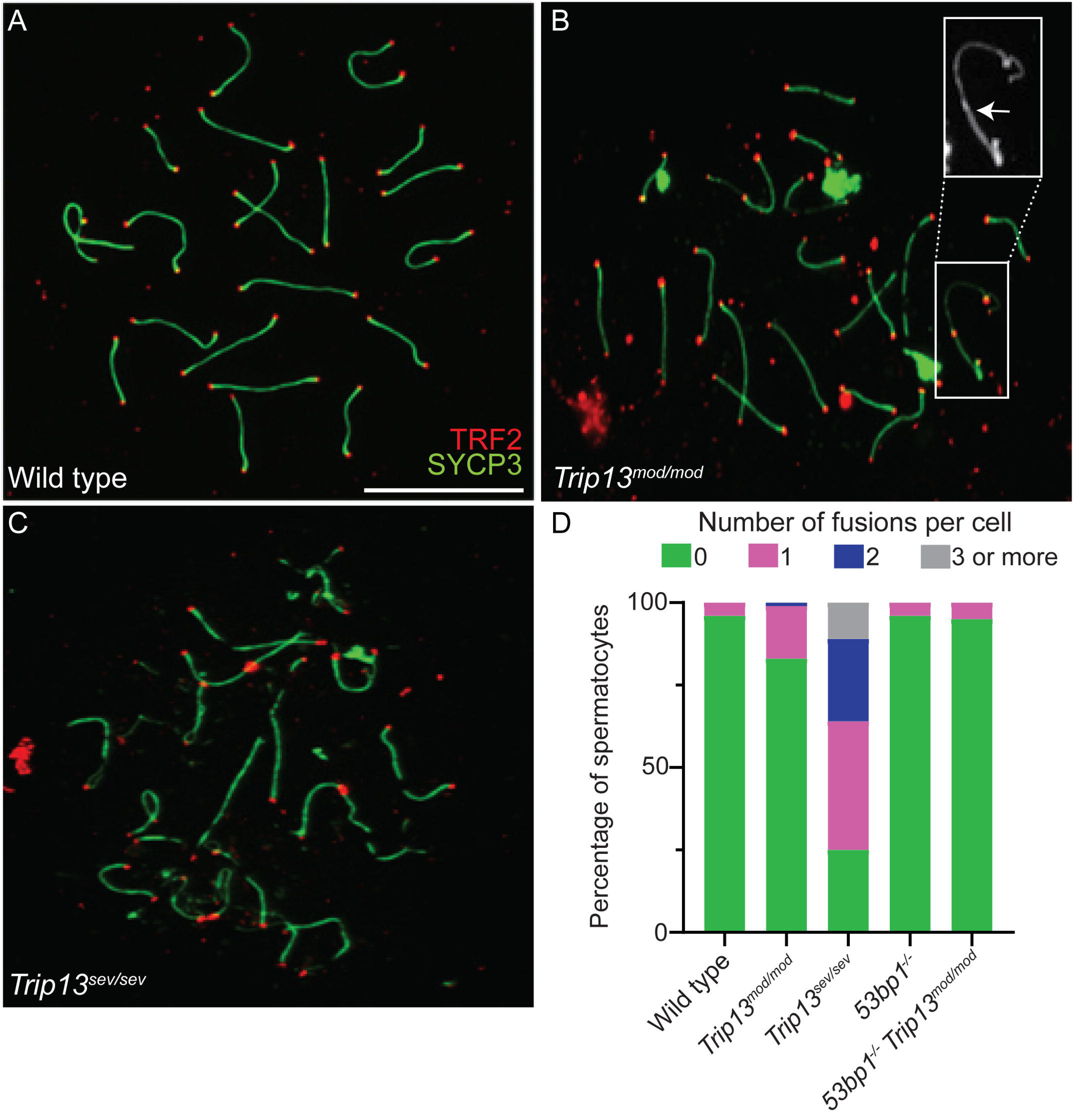
TRIP13 promotes DSB repair by homologous recombination during meiotic prophase. (A,B) Representative images of pachytene-like *Dmc1^-/-^ Chk2^-/-^* (A) and *Dmc1^-/-^ Chk2^-/-^ Trip13^mod/mod^* (B) spermatocytes stained for SYCP3 and the DNA damage marker γH2AX. (C) Quantification of γH2AX patches per cell. The line denotes the mean. Each dot represents a cell. The mean, SD, and the number of cells analyzed (N) are shown (D-L). Two animals were used per each genotype for these analysis. Displayed as in panels A–B, representative images and quantification of RAD51 foci (D–F), RPA foci (G–I), and REV7 foci (J–L). P values are from t tests.

During meiotic prophase, DSBs are preferentially repaired by a meiosis-specific version of homologous recombination that, among other particularities, preferentially uses the homologous chromosome as the template, instead of the sister chromatid that is preferred in somatic cells (Huang and Roig, 2023). Other pathways, like NHEJ or somatic-like (intersister) homologous recombination, are restricted to later stages of meiotic prophase when homologous pairing and exchange of DNA between homologous chromosomes have already occurred (Enguita-Marruedo et al., 2019).

To gain insight into the fate of DSBs in *Dmc1^-/-^ Chk2^-/-^ Trip13^mod/mod^* triple mutant spermatocytes, we analyzed chromosome-associated foci of the RAD51 recombinase, a marker for recombination. We found that triple mutant cells had fewer foci than double mutants (Fig. 5D-F). Similar results were obtained for the ssDNA binding protein RPA, which labels resected DSBs and ssDNA from D-loops (Fig. 5G-I). These data suggest that the number of DSBs being repaired by meiotic recombination in *Dmc1^-/-^ Chk2^-/-^ Trip13^mod/mod^* cells is significantly reduced compared to *Dmc1^-/-^ Chk2^-/-^* cells, which agrees with a possible role for TRIP13 in the regulation of DSB repair pathway choice during meiosis.

In contrast to the markers of recombination, REV7 focus numbers were elevated in the triple mutant spermatocytes as compared to *Dmc1^-/-^ Chk2^-/-^* double mutants (Fig. 5J-L). This finding is consistent with TRIP13 inhibiting the formation of DSB-associated Shieldin complexes, and thus inhibiting NHEJ, during meiosis. Importantly, in both control and *Trip13* mutant cells, the number of REV7 foci were significantly lower than the levels observed for recombination markers (Fig. 5).

### TRIP13 prevents 53BP1-dependent chromosome end fusions during meiotic prophase

To gather further evidence that TRIP13 inhibits NHEJ during meiotic prophase I, we asked whether there is an increased frequency of chromosomal end-to-end fusions in *Trip13* single mutant spermatocytes. Such fusions were rare in wild-type spermatocyte spreads, affecting less than 0.1% of chromosome ends and being present in fewer than 4% of spermatocytes (Fig. 6A, D). However, fusions were more frequent in both *Trip13* mutants, increasing to 0.36% of chromosome ends and 16.8% of *Trip13^mod/mod^* spermatocytes (P = 0.0023, G test) and increasing even more to 3.27% of chromsome ends and 75% of *Trip13^sev/sev^* spermatocytes (P=9.7E-16 compared to control and P=1.21E-11 compared to *Trip13^mod/mod^*, G test) (Fig. 6B-D). The increase in end fusions caused by TRIP13 defiency appears to be entirely dependent on 53BP1, as the *Trip13^mod^* mutation did not trigger more fusions in a *53bp1^-/-^* background (P=0.791, t test, Fig. 6D). This 53BP1 dependence indicates that the end fusions in *Trip13* mutant cells occurred through NHEJ, in turn implying that TRIP13 regulates DSB repair pathway choice in meiotic cells.

**Figure 6.**
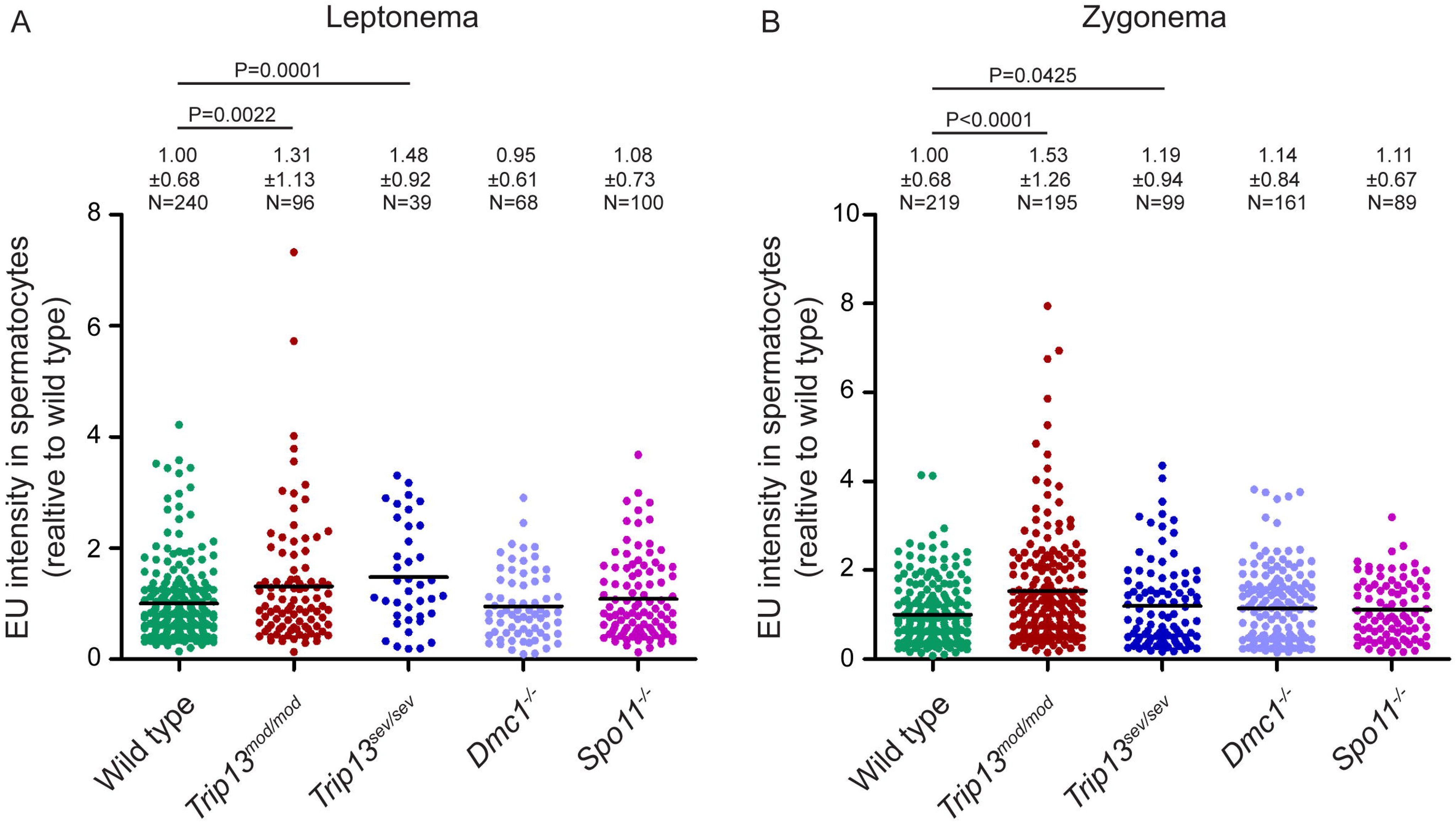
TRIP13 prevents chromosome end fusions. (A-C) Representative images of wild type (A), *Trip13^mod/mod^* (B) and *Trip13^sev/sev^* (C) spermatocyte spreads stained for SYCP3 and the telomeric protein TRF2. The scale bar represents 10 μm and applies to all images. The arrow highlights a chromosome end fusion. (D) Proportion of cells displaying the indicated numbers of chromosome end fusions. Number of animals used for this analysis: WT: 4 mice; *Trip13^mod/mod^*: 3 mice; *Trip13^sev/sev^*: 2 mice; *53bp1^-/-^*: 3 mice; *53bp1^-/-^Trip13^mod/mod^*: 3 mice.

## Discussion

TRIP13 is a multifunctional AAA+ ATPase that orchestrates diverse aspects of somatic and meiotic cell processes in mammals, such as chromosome segregation, DSB repair, chromosome axis remodeling, completion of homologous synapsis, regulation of CO formation, and silencing of the XY chromosomes (Roig et al., 2010; Nelson et al., 2015; Pacheco et al., 2015; Clairmont et al., 2020). Our study reveals two novel roles for TRIP13 in regulating transcriptional activity and DNA repair pathway choice in meiotic cells. These findings expand our understanding of TRIP13 beyond its previously described functions.

We found *Trip13* mutant cells had higher levels of EU incorporation in meiotic prophase, corroborated by finding more pRNAP on chromatin. This increased transcription can not be attributed to a meiotic arrest, as *Dmc1* and *Spo11* mutants did not display a similar phenotype. Notably, the findings also suggest that meiotic transcription repression during prophase I operates as part of a meiotic program that is independent of DSB formation or repair completion. Our findings are in line with a recent study that suggests that this transcription inhibition is dependent on *Stra8* and initiates before the meiotic onset (Bellutti et al., 2024).

Our RNAseq analysis revealed a pronounced upregulation of genes located on the X and Y chromosomes in *Trip13* mutant samples, highlighting a critical role for TRIP13 in the regulation of sex chromosome gene expression. This finding aligns with our previous studies demonstrating that TRIP13 is essential for the proper recruitment of ATR to the unsynapsed axes of the sex chromosomes, thereby facilitating ATR-mediated chromatin signaling (Pacheco et al., 2015; Marcet-Ortega et al., 2017). This mechanism is likely driven by the persistence of HORMAD1 on synapsed chromosomes in *Trip13* mutants (Wojtasz et al., 2009; Roig et al., 2010). The widespread retention of HORMAD1 mimics an unsynapsed signal, which interferes with the correct assembly of the MSCI machinery and may ultimately lead to MSCI failure (Mahadevaiah et al., 2008). These observations suggest that TRIP13 contributes to MSCI by regulating the chromosomal axis through the timely removal of HORMAD1 as synapsis progresses. Indeed, expression analysis performed in 14-day-old testis from HORMAD1 mutants revealed that around 70% of the top upregulated genes mapped on the sex chromosomes (Shin et al., 2010).

Notably, approximately one-third of the upregulated genes in *Trip13* mutants were located on autosomes, indicating that the transcriptional dysregulation extends beyond the sex chromosomes. This broader transcriptional activation suggests that TRIP13 may also play a role in repressing gene expression at the onset of meiosis. The molecular basis of this potential function remains unclear, but it may similarly involve HORMAD1 remodeling, as *Hormad1^-/-^* mice also present a similar phenotype (Shin et al., 2010). Further investigation will be necessary to elucidate whether TRIP13’s role in transcriptional repression is mechanistically linked to its regulation of HORMAD1 dynamics.

Given that MSCI is a specialized form of MSUC, and having established TRIP13’s role in facilitating sex chromosome silencing in spermatocytes, we next investigated its potential involvement in MSUC-mediated oocyte arrest (Cloutier et al., 2015). In mice, oocyte development can be halted in response to defects in homologous synapsis or incomplete recombination (Hunter, 2017; Huang and Roig, 2023). Previous studies have demonstrated that the arrest of synapsis-deficient oocytes relies on the same molecular machinery required for MSCI (Daniel et al., 2011; Wojtasz et al., 2012; Cloutier et al., 2015; Hunter, 2017).

Our data suggest that TRIP13 does not play a central role in MSUC, as *Trip13* mutation only partially rescues oocyte numbers in one of the two tested models of synapsis failure. Specifically, *Trip13* mutation restored oocyte numbers in *Dmc1^-/-^ Chk2^-/-^* mice, but not in *Spo11^-/-^* mice. Although both models exhibit synapsis defects, *Dmc1^-/-^ Chk2^-/-^* mutants also accumulate unrepaired DSBs, which activate a CHK1-dependent DNA damage response that contributes to oocyte elimination (Martínez-Marchal et al., 2020; Rinaldi et al., 2020). Therefore, the rescue observed in *Dmc1^-/-^ Chk2^-/-^* mutants likely reflects enhanced repair of DSBs rather than a direct effect on synapsis-related arrest.

Direct assessment of the hypothesis that reduced TRIP13 activity in *Dmc1^-/-^ Chk2^-/-^* oocytes may facilitate DSB repair was limited by scarcity of oocyte samples. Nonetheless, our analysis in spermatocytes supports this model, as *Trip3* reduction in *Dmc1^-/-^ Chk2^-/-^* mutant cells correlates with a decrease in markers of unrepaired DSBs, suggesting improved repair efficiency.

The observed reduction in recombination markers such as RPA and RAD51 in *Trip13* mutants suggests a decreased engagement of homologous recombination in DSB repair during meiotic prophase. Alternatively, repair by recombination using the sister chromatid as a template might proceed more efficiently in this background, despite the persistence on chromosomes of HORMAD1, which is thought to help enforce preferential use of the homolog (Humphryes and Hochwagen, 2014).

The increased number of REV7 foci in these mutants implies that TRIP13 may regulate REV7 dynamics in meiosis, similar to its role in somatic cells (Clairmont et al., 2020; Sarangi et al., 2020). These findings are consistent with the hypothesis that, in the absence of TRIP13, at least some DSBs may be redirected towards NHEJ, a pathway typically suppressed during meiotic prophase I (Enguita-Marruedo et al., 2019).

However, the total number of REV7 foci in *Dmc1^-/-^ Chk2^-/-^ Trip13^mod/mod^* cells was modest compared to the expected number of DSBs or recombination events. This discrepancy may reflect differences in the turnover rates of homologous recombination and NHEJ markers, as NHEJ is known to resolve DSBs more rapidly than homologous recombination (Mao et al., 2008). Alternatively, it is possible that NHEJ-mediated repair in *Trip13* mutants accounts for only a small fraction of DSBs. Further studies are needed to clarify the mechanistic role of TRIP13 in regulating DSB repair during meiotic prophase.

Our findings suggest that the mutation of *Trip13* in a *Dmc1* mutant background causes repair to occur predominantly through end-joining pathways rather than homologous recombination. This is particularly intriguing given that Dmc1 mutants are known to exhibit extensive hyper-resection of DSB ends, which typically favors HR (Paiano et al., 2020; Yamada et al., 2020). When *Trip13* expression is lowered, however, these long single-stranded regions may become substrates for alternative end-joining mechanisms. One plausible explanation is the engagement of the deletion-prone microhomology-mediated end joining (MMEJ), a pathway that exploits exposed microhomologies created by hyper-resection (Kumari et al., 2025). The increased REV7 signal, associated with Shieldin complex activity, further supports a model in which TRIP13 normally suppresses inappropriate end-joining during meiosis. Thus, the combined phenotype of fewer unrepaired breaks, reduced homologous recombination markers, and elevated pro-NHEJ factors points to a compensatory shift toward error-prone MMEJ, which may rescue repair at the cost of genomic integrity.

Finally, our findings undescore a critical role for TRIP13 in preventing chromosome end fusions mediated by end joining mechanisms. Previous studies have shown that TRIP13 localizes to chromosome ends during meiotic prophase I (Gómez-H et al., 2019; Chotiner et al., 2024). This localization appears to be specific to mammals, as TRIP13 orthologs have not been observed at chromosome ends in other model organisms (San-Segundo and Roeder, 1999; Joyce and McKim, 2010; Deshong et al., 2014; Lambing et al., 2015). Our data suggest that TRIP13 plays a protective role in maintaining genome integrity by preventing inappropriate DSB repair via NHEJ at chromosome ends, thereby avoiding deleterious chromosome fusions.

In somatic cells, telomeres are safeguarded from NHEJ-mediated fusions by shelterin complexes (de Lange, 2018). However, during meiosis, shelterin is partially replaced by a meiosis-specific complex composed of TERB1/2-MAJIN (Shibuya et al., 2015), which may render telomeres more susceptible to aberrant repair. This meiosis-specific teleromere complex is essential for proper chromosome dynamics, including homolog pairing, synapsis, and recombination. Thus, TRIP13 may serve as an additional layer of protection against an intrinsic vulnerability of chromosome ends during meiotic prophase.

Collectively, our data strongly support a model in which TRIP13 regulates DSB repair pathway choice during meiosis, promoting homologous recombination over NHEJ. Importantly, even in the complete absence of TRIP13, homologous recombination-mediated repair can still occur, as evidenced by the successful homolog pairing observed in *Trip13^sev/sev^* and *Trip13* knockout mice (Roig et al., 2010; Chotiner et al., 2024). This suggests that additional mechanisms contribute to the regulation of DSB repair pathway selection during meiosis.

In summary, our study reveals new and multifaceted roles for the TRIP13 protein, positioning it as a key regulator of early meiotic events. By orchestrating DSB repair pathway choice, protecting chromosome ends, and facilitating essential processes such as meiotic silencing, transcriptional regulation, and recombination, TRIP13 emerges as a central player in preserving genomic integrity during gametogenesis. Future research should aim to determine whether these meiotic functions of TRIP13 are conserved across diverse species, which could provide broader evolutionary insights into its role in genome stability.

## Acknowledgments

This work was supported by Spanish Ministerio de Ciencia, Innovación e Universidades grants (I.R.: PID 2019-107082RB-I00, PID2022-138905OB-I00) and the US National Institutes of Health (S.K.: R35 GM118092; M.J.: R01HD112624). M.M.-O. was the recipient of an FPI fellowship from the Spanish Ministerio de Ciencia, Innovación e Universidades (BES-2011-045381)), and travel fellowships EEBB-I-14-08517 and EEBB-I-15-09556 from the Spanish Ministerio de Economía y Competitividad. MSK core facilities are supported by the US National Cancer Institute cancer center support grant P30 CA08748. Institutional support to CNAG was provided by the Spanish Ministry of Science and Innovation through the Instituto de Salud Carlos III, and by the Generalitat de Catalunya through the Departament de Salut and the Departament de Recerca i Universitats. We would like to acknowledge the rest of the members of the Roig Lab and the Keeney Lab for their support, comments, and discussions about the project.

## Material and Methods

### Experimental mice and genotyping

Mice carrying *Trip13*, *Chk2, Spo11* and *Dmc1* mutations were previously generated and described elsewhere (Yoshida et al., 1998; Baudat et al., 2000; Takai et al., 2002; Roig et al., 2010). These lines were maintained in a C57Bl/6-129/Sv mixed background. All experiments were performed using at least three animals (unless mentioned in the text) and compared with control littermates when possible, or from animals of closely related parents. The term wild-type (WT) in the text and figures refers to both homozygous and heterozygous mice.

Experiments performed in this study comply with US and EU regulations and were approved by the Ethics Committee of the UAB and Catalan Government and by the MSKCC Institutional Animal Care and Use Committee.

Mouse genotyping was performed by PCR analysis from the DNA extracted from the tails as previously performed (Yoshida et al., 1998; Baudat et al., 2000; Takai et al., 2002; Roig et al., 2010; Marcet-Ortega et al., 2017).

### Testis and Ovaries collection

Mice were euthanized using CO2 asphyxiation followed by cervical dislocation to ensure death. Testes or ovaries were immediately excised, cleaned of surrounding adipose tissue, and washed in cold phosphate-buffered saline (PBS). Next, samples were treated differently based on their used.

### RNA Extraction, Quantification and Sequencing

Total RNA was extracted from the testis tissue using the RNeasy Plus mini kit (Qiagen) according manufacturer’s instructions.

RNA concentration and purity were assessed using a NanoDrop 2000 spectrophotometer (Thermo Scientific). The integrity of RNA was evaluated with the 2100 Bioanalyzer (Agilent).

RNA libraries were prepared using the TruSeq Stranded mRNA Library Prep Kit (Illumina) following the manufacturer’s instructions. The libraries were quantified using the Qubit 2.0 Fluorometer (Life Technologies) and their size distribution was assessed using the Agilent 2100 Bioanalyzer. The libraries were then pooled in equimolar concentrations and sequenced on an Illumina platform (Illumina) to generate 75 bp paired-end reads.

The pair-end reads were mapped with the software GEM-Spit-RNA-Mapper (to map the reads to the mouse genome reference annotation GRCm38) (Marco-Sola et al., 2012). To quantify the transcripts that maps to each gene Flux Capacitator software was used (Montgomery et al., 2010). Then, to study the differential expression, and to normalize the number of reads per gene/total of mapped reads, between the wild type and mutant mice the R EdgeR software was used (Robinson et al., 2010). This algorithm normalizes the data with the TMM method (Robinson and Oshlack, 2010) considering different library sizes and different RNA composition between the samples.

To validate the RNA sequencing results, quantitative reverse transcription PCR (qRT-PCR) was performed on selected differentially expressed genes. cDNA was synthesized from 2 μg of total RNA using the iScript Advanced cDNA Synthesis (Bio-Rad). qRT-PCR was conducted using iTaq Universal SYBR Green Supermix (Bio-Rad) and intron-spanning PrimePCR primers (Bio-Rad) on CFX96™ Real-Time PCR Detection System thermocycler (Bio-Rad) and analyzed with Bio-Rad CFX Manager 3.1 software.

All sequencing data have been deposited in the NCBI Gene Expression Omnibus (GEO) under the accession number [submission pending].

### Histology and staining

Fresh mouse tissues were fixed overnight at 4℃ with Bouin’s fixative. After fixation, tissues were washed in PBS, dehydrated in a series of ethanol with increasing concentration, cleared with histoclear and infiltrated with paraffin in sequence.

Embedded tissues in paraffin blocks were cut into 5-7 µm slices using a microtome and dried overnight at 37℃. Before staining with PAS-Hematoxylin, slides were deparaffinized in Xylene and rehydrated in a series of ethanol with decreasing concentration.

### Nuclei spreading and immunofluorescence

Spermatocyte nuclei spreading was prepared using frozen testes. Protocol was adapted from (Roig et al., 2004): briefly, a small portion of testis was cut and minced thoroughly with a sterile blade in cold PBS (pH 7.4) containing 1× protease inhibitor, PI (Roche Diagnostics) on a petri dish; cell mixture was transferred to a sterile Eppendorf and sat for 15 min to sediment; 25 µl of supernatant cell suspension was spread onto a glass slide and incubated with 1% Lipsol containing 1× PI, followed by fixation in PFA fixative solution 2 h at room temperature in a closed humid chamber. Slides were dried under a fume hood and then washed in 0.4% Photoflo (Kodak).

For immunofluorescence staining, slides were blocked in 0.2% BSA, 0.2% gelatin, 0.05% Tween-20 in PBS at room temperature, followed by incubation of primary antibody diluted in blocking solution in a humid chamber overnight at 4℃. The next day, slides were washed four times in blocking solution, and incubated in secondary antibody diluted in blocking solution in a humid chamber for 1 h at 37℃, followed by another four washes in blocking solutions. Drained slides were mounted with 0.1 µg/ml DAPI in Vectashield antifade mounting medium. and analyzed with an epifluorescence microscope (Zeiss Axioskop). Primary antibodies used for IF: anti-phospho(S2)-RNA pol II Ms(abcam, 1:100), anti-SYCP3 Ms or Rb (abcam, 1:200), anti-phospho-Histone H2A.X Ms (Millipore, 1:400), anti-RPA32 (4E4) Rat (cell signaling, 1:100), anti-RAD51 (ab-1) Rb (Millipore, 1:100), and anti-REV7 Rb (abcam, 1:100).

### Follicle count and classification

The whole ovaries were sectioned for follicle quantification. Every fifth section of each ovary was counted to assess oocyte counts and follicle proportions. The oocytes were counted manually under the bright field Zeiss Axioskop microscope only if the nucleus was clearly visible. Each oocyte was classified in primordial follicles (with one layer of flat granulosa cells), primary follicles (with one layer of cuboid granulosa cells), secondary follicles (with two or more layers of granulosa cells) and antral follicles (with the antrum) and abnormal follicles (big oocytes with one layer or without granulosa cells).

### EU Imaging on spermatocytes

To analyze global RNA expression in meiotic cells, we used the EU Click-it Imaging technique (Click-iT® RNA Alexa Fluor® 488 Imaging Kit, Life Technologies) to cytologically detect newly synthesized RNA. Briefly, we enzymatically dissociated mouse testis to obtained a cell suspension, as previously performed (Cole et al., 2014). Briefly, testis were decapsulated and seminiferous tubules were incubated in GBSS with 0.5 mg/ml collagenase (Sigma) for 15 min shaking vigorously every five minutes. The tubules were rinsed with GBSS and incubated with 0.5 mg/ml trypsin (Sigma) supplemented with 2 μg/ml DNase I (Sigma) 15 min. Trypsin was inactivated by addition of fetal bovine serum. The solution was filtered through a 70-μM cell strainer (BD Falcon) to obtain a cell suspension. Then, the cell suspension was incubated for 2.5h in 5 mM 5-Ethynyl Uridine (EU) in of RPMI medium (containing 10% FCS and 1x Penicillin/Spretomycin/Fungizone).

After this time, slides were prepared as previously done (Mahadevaiah et al., 2009; Pacheco et al., 2015; Marcet-Ortega et al., 2017). Briefly, 60 μl of cell suspension were placed on Superforst slides over a cold platform. 10 minutes later, four drops of CSK buffer were added and slides were incubated for 13 minutes. Then, six drops of 4% paraformaldehyde in PBS were added. Slides were incubated for 30 minutes.

After this time, the cold platform was tilted to eliminate excess of liquid from the slides. Slides were then rinsed in cold PBS and air dried.

Then, the modified RNA was labeled with Alexa Fluor 488 through a chemoselective ligation reaction (Click-it^TM^, Invitrogene) according to the manufacturer’s instructions. Slides were washed two times in cold PBS, then washed with cold 0.5% Triton X-100 in PBS for 30 minutes to permeabilize the cells. After washing the slides in cold PBS, 100 μl of the Click-iT Reaction master mix were added to each slide and incubated in a humid chamber for 30 min. Slides were washed with Rinse buffer and PBS. Then, slides were processed for immunostaining, first incubating them in blocking solution (0.2% BSA, 0.2% gelatin, 0.05% Tween-20 in PBS) for 10 minutes. Then we added the primary antibodies in PTBG with 2 mM Vanadyl ribonucleoside: Streptavidin Alexa Fluor 488 (1:100, Life Technologies), mouse anti-SYCP3 at 1:100 (Santa Cruz) and rabbit anti-γH2AX (1:100, Millipore) and incubated it for 1 h at 37°C. Slides were washed three times in PBS and secondary antibodies diluted in PTBG with 2 mM Vanadyl ribonucleoside were added (anti-mouse Alexa Fluor 594 (1:200) and anti-rabbit Alexa Fluor 647 (1:200)). Slides were then washed three times with PBS and mounted with Vectashield mounting medium with DAPI at a concentration of 1.5 μg/ml. Finally, slides were visualized under a fluorescence microscope.

For phosphorylated RNA polymerase II analysis, slides were prepared using this protocol but cells were incubated without EU. Slides were blocked in blocking solution, followed by incubation primary antibody in a humid chamber overnight at 4℃. The primary antobodies used were: mouse anti-phospho(S2)-RNA pol II Ms (abcam, 1:100), goat anti-SYCP3 (1:100, Santa Cruz) and rabbit anti-γH2AX (1:100, Santa Cruz). Slides were washed in blocking solution, and incubated in secondary antibody for 1 hour at 37℃, followed by another four washes in blocking solution.

Secondary antibodies used were: anti-mouse Alexa Fluor 488, anti-goat Alexa Fluor 594 (1:200) and anti-rabbit Alexa Fluor 647 (1:200) from Molecular Probes.

EU incubated spermatocytes or RNApol II stained slides were visualized in a Zeiss Axio2 Observer inverted epifluorescence microscope using a 63x oil immersion objective and images were captured with a CCD camera. Images were acquired at 0.5 μm stacks interval from the top to the bottom of the cell. Images were then processed with Slidebook software package (Intelligent Imaging Innovations) to create a sum projection image from the stacks. To measure fluorescence intensity we used ImageJ 1.49g software (National Institutes of Health, USA; http://imagej.nih.gov/ij/). EU intensity signal was quantified by taking into account the nuclei area identified with DAPI, and obtained background signal from the surrounding areas. We used ImageJ parameter Integrated density to plot the graphics (which is the sum of the values of the pixels in that selection, equivalent to the product of the Area and the Mean Gray Value –fluorescence intensity values-).

### Image processing and data analysis

Microscopy analysis was performed with a Zeiss Axiophot microscope. Images were captured with a Point Gray Research, Inc. camera with the ACO XY Software (A. COLOMA Open microscopy). All images were processed with Adobe Photoshop CC to overlay the different fluorescent channels and match the intensity observed in the microscope.

### Statistical analysis

Data analysis and statistical inference were performed using the GraphPad Prism 8 software (https://www.graphpad.com/scientific-software/prism/).

---

**Figure S1.**
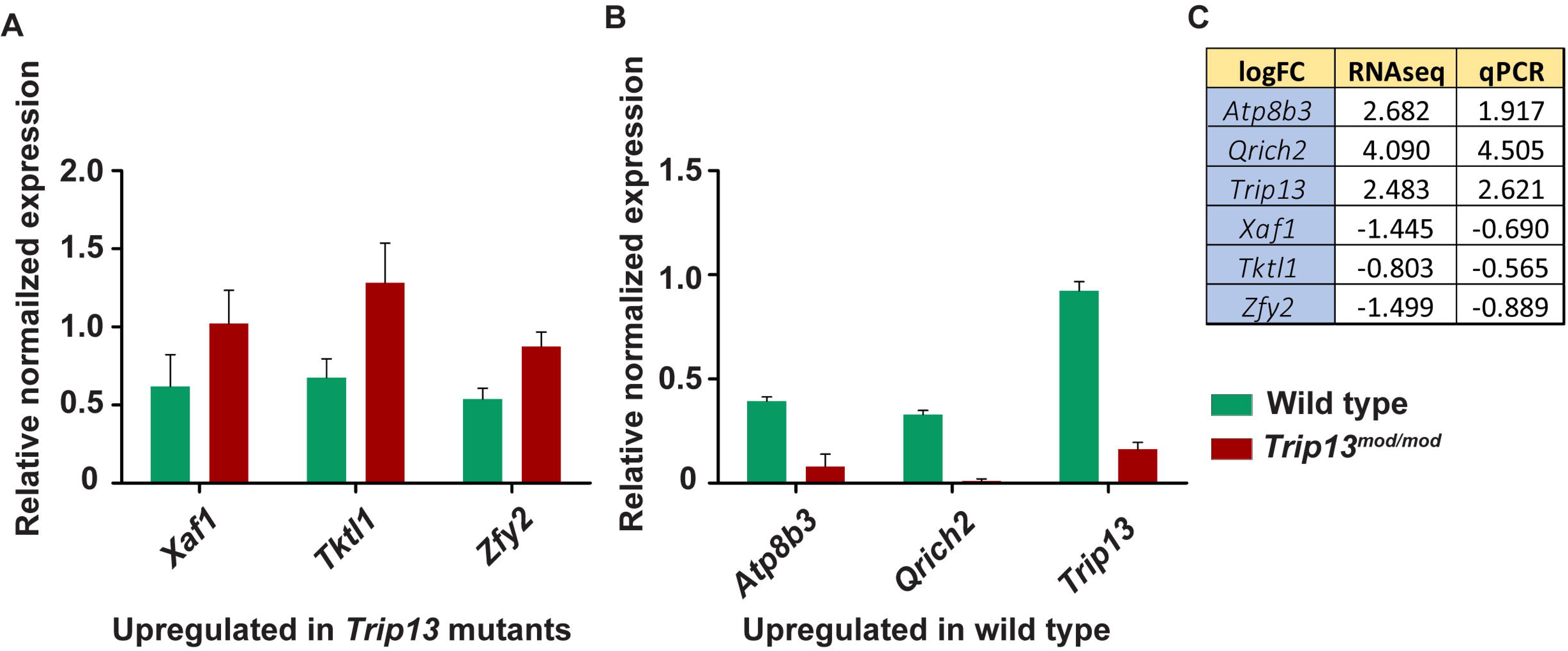
Comparison of global RNA synthesis in *Trip13^mod/mod^*, *Trip13^sev/sev^*, *Dmc1^-/-^* and *Spo11^-/-^* mutants. Quantification of the EU intensity in leptotene (A) and zygotene (B) spermatocytes of the indicated genotypes relative to the average in wild type. Horizontal lines represent means. Means ± SD and the number of counted cells (N) are indicated. P values are from t tests comparing to wild type. Number of mice analyzed per stage and genotype: Leptonema: WT:13 mice, *Trip13^mod/mod^*: 5 mice, *Trip13^sev/sev^*: 1 mouse, *Dmc1^-/^*: 3 mice, *Spo11^-/-^*: 3 mice. Zygonema: WT:10 mice, *Trip13^mod/mod^*: 5 mice, *Trip13^sev/sev^*: 1 mouse, *Dmc1^-/^*: 4 mice, *Spo11^-/-^*: 3 mice.

**Figure S2.**
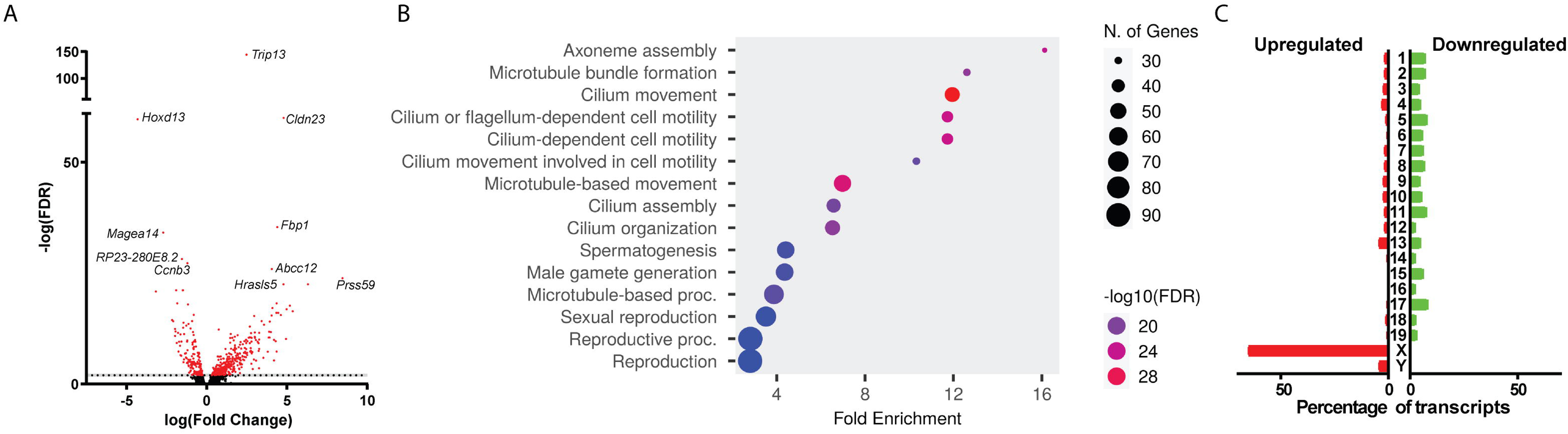
Validation of RNA sequencing results by real-time PCR. (A,B) Normalized expression relative to housekeeping gene *Hmbs* in the wild type (green) and *Trip13^mod/mod^*(red) of genes upregulated (A) or downregulated (B) in *Trip13* mutants. Error bars represent mean ± SD of N = 3 mice per genotype. (C) Differential expression relative to wild type expressed as log_2_ fold change (logFC) from RNA sequencing (RNAseq) and real-time PCR (qPCR).

**Table S1.**
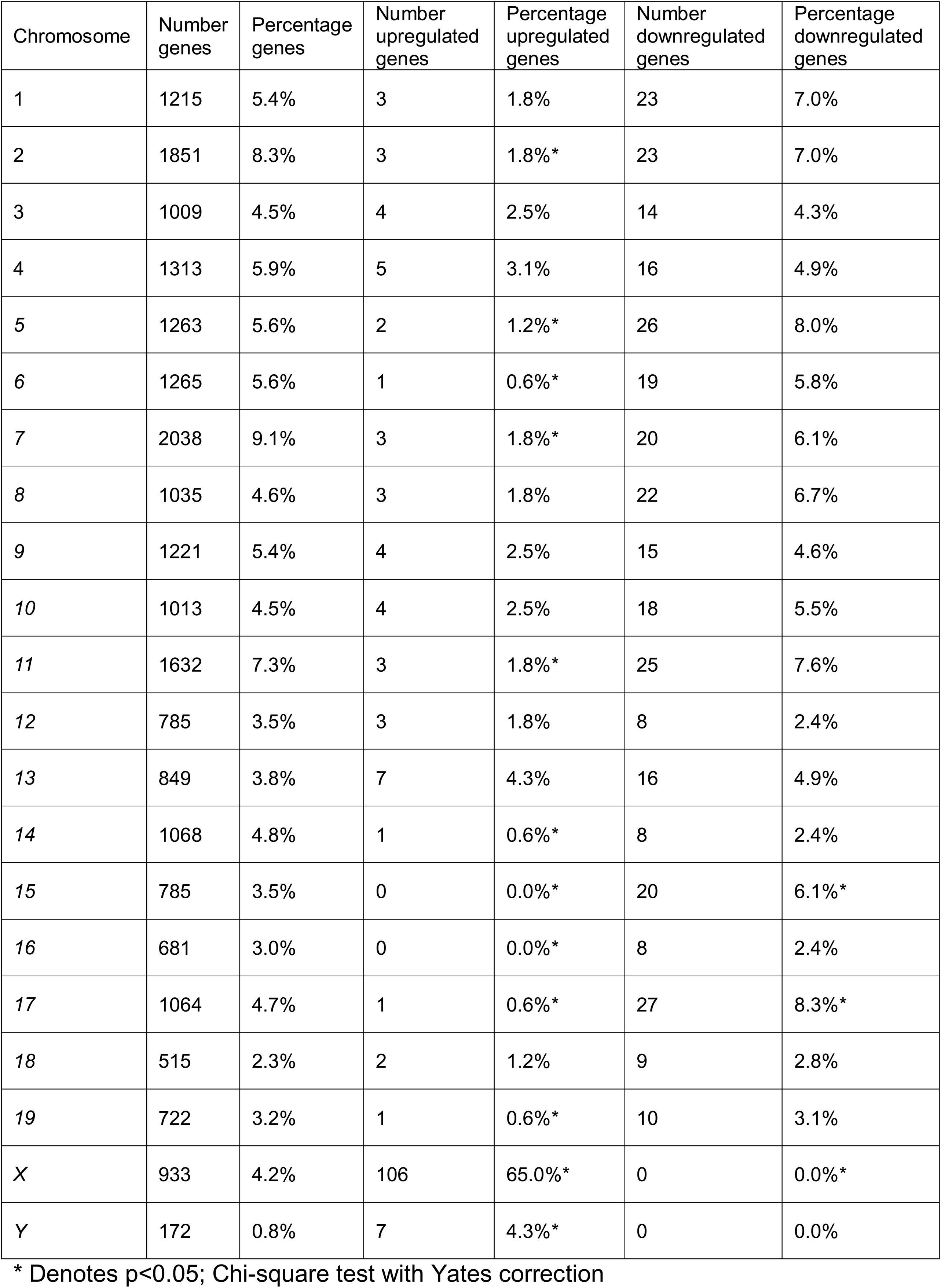
Correlation of the number of genes per chromosome and the number of upregulated and downregulated genes in *Trip13^mod/mod^* testis.

**Table S2.**
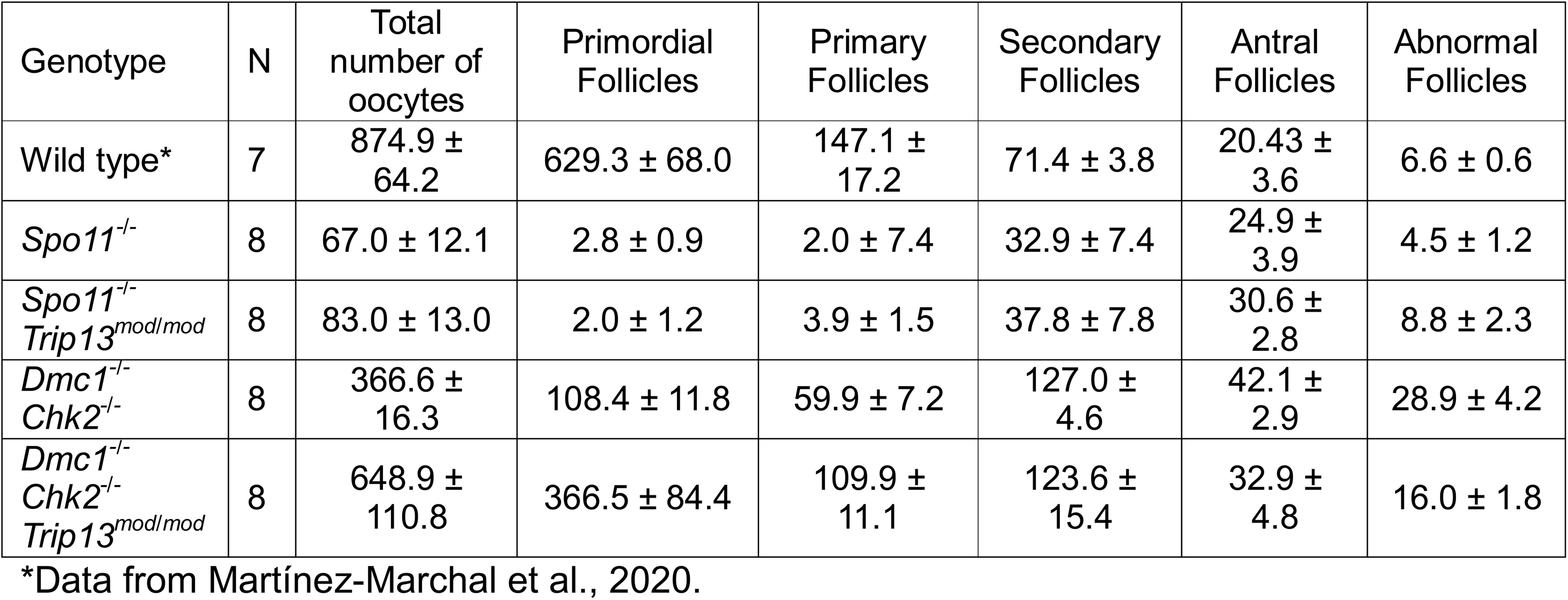
Oocyte number and follicle type in 1-month-old mouse ovaries. The numbers express the average ± SEM. N indicates the number of ovaries counted.

| Genotype | N | Total number of oocytes | Primordial Follicles | Primary Follicles | Secondary Follicles | Antral Follicles | Abnormal Follicles |
| --- | --- | --- | --- | --- | --- | --- | --- |
| Wild type* | 7 | 874.9 ± 64.2 | 629.3 ± 68.0 | 147.1 ± 17.2 | 71.4 ± 3.8 | 20.43 ± 3.6 | 6.6 ± 0.6 |
| *Spo11*<sup>-/-</sup> | 8 | 67.0 ± 12.1 | 2.8 ± 0.9 | 2.0 ± 7.4 | 32.9 ± 7.4 | 24.9 ± 3.9 | 4.5 ± 1.2 |
| *Spo11*<sup>-/-</sup><br>*Trip13*<sup>mod/mod</sup> | 8 | 83.0 ± 13.0 | 2.0 ± 1.2 | 3.9 ± 1.5 | 37.8 ± 7.8 | 30.6 ± 2.8 | 8.8 ± 2.3 |
| *Dmc1*<sup>-/-</sup><br>*Chk2*<sup>-/-</sup> | 8 | 366.6 ± 16.3 | 108.4 ± 11.8 | 59.9 ± 7.2 | 127.0 ± 4.6 | 42.1 ± 2.9 | 28.9 ± 4.2 |
| *Dmc1*<sup>-/-</sup><br>*Chk2*<sup>-/-</sup><br>*Trip13*<sup>mod/mod</sup> | 8 | 648.9 ± 110.8 | 366.5 ± 84.4 | 109.9 ± 11.1 | 123.6 ± 15.4 | 32.9 ± 4.8 | 16.0 ± 1.8 |

\*Data from Martínez-Marchal et al., 2020.

## References

Anderson, E. L., Baltus, A. E., Roepers-Gajadien, H. L., Hassold, T. J., de Rooij, D. G., van Pelt, A. M. M., et al. (2008). *Stra8* and its inducer, retinoic acid, regulate meiotic initiation in both spermatogenesis and oogenesis in mice. Proceedings of the National Academy of Sciences 105, 14976–14980. doi: 10.1073/pnas.0807297105

Baudat, F., Manova, K., Yuen, J. P., Jasin, M., and Keeney, S. (2000). Chromosome synapsis defects and sexually dimorphic meiotic progression in mice lacking Spo11. Mol Cell 6, 989–998. Available at: http://www.ncbi.nlm.nih.gov/entrez/query.fcgi?cmd=Retrieve&db=PubMed&dopt=Citation&list_uids=11106739

Bellutti, L., Pen, E. C. S., Guerquin, M.-J., Messiaen, S., Llano, E., Cluzet, V., et al. (2024). Genome-wide transcriptional silencing and mRNA stabilization allow the coordinated expression of the meiotic program in mice. bioRxiv, 2024.02.08.579523. doi: 10.1101/2024.02.08.579523

Bolcun-Filas, E., and Handel, M. A. (2018). Meiosis: the chromosomal foundation of reproduction. Biol Reprod 99, 112–126. doi: 10.1093/biolre/ioy021

Bolcun-Filas, E., Rinaldi, V. D., White, M. E., and Schimenti, J. C. (2014). Reversal of Female Infertility by Chk2 Ablation Reveals the Oocyte DNA Damage Checkpoint Pathway. Science (1979) 343, 533–536. doi: 10.1126/science.1247671

Burgoyne, P. S., Mahadevaiah, S. K., and Turner, J. M. (2009). The consequences of asynapsis for mammalian meiosis. Nat Rev Genet 10, 207–216. Available at: http://www.ncbi.nlm.nih.gov/entrez/query.fcgi?cmd=Retrieve&db=PubMed&dopt=Citation&list_uids=19188923

Chotiner, J. Y., Leu, N. A., Yang, F., Cossu, I. G., Guan, Y., Lin, H., et al. (2024). TRIP13 localizes to synapsed chromosomes and functions as a dosage-sensitive regulator of meiosis. Elife 12. doi: 10.7554/eLife.92195

Clairmont, C. S., Sarangi, P., Ponnienselvan, K., Galli, L. D., Csete, I., Moreau, L., et al. (2020). TRIP13 regulates DNA repair pathway choice through REV7 conformational change. Nat Cell Biol 22, 87–96. doi: 10.1038/s41556-019-0442-y

Cloutier, J. M., Mahadevaiah, S. K., ElInati, E., Tóth, A., and Turner, J. (2015). Mammalian meiotic silencing exhibits sexually dimorphic features. Chromosoma. doi: 10.1007/s00412-015-0568-z

Cole, F., Baudat, F., Grey, C., Keeney, S., de Massy, B., and Jasin, M. (2014). Mouse tetrad analysis provides insights into recombination mechanisms and hotspot evolutionary dynamics. Nat Genet 46, 1072–1080. doi: 10.1038/ng.3068

Daniel, K., Lange, J., Hached, K., Fu, J., Anastassiadis, K., Roig, I., et al. (2011). Meiotic homologue alignment and its quality surveillance are controlled by mouse HORMAD1. Nat Cell Biol 13, 599–610. doi: 10.1038/ncb2213

de Lange, T. (2018). Shelterin-Mediated Telomere Protection. Annu Rev Genet 52, 223–247. doi: 10.1146/annurev-genet-032918-021921

Deshong, A. J., Ye, A. L., Lamelza, P., and Bhalla, N. (2014). A quality control mechanism coordinates meiotic prophase events to promote crossover assurance. PLoS Genet 10, e1004291. doi: 10.1371/journal.pgen.1004291

Enguita-Marruedo, A., Martín-Ruiz, M., García, E., Gil-Fernández, A., Parra, M. T., Viera, A., et al. (2019). Transition from a meiotic to a somatic-like DNA damage response during the pachytene stage in mouse meiosis. PLoS Genet 15. doi: 10.1371/journal.pgen.1007439

Ghezraoui, H., Oliveira, C., Becker, J. R., Bilham, K., Moralli, D., Anzilotti, C., et al. (2018). 53BP1 cooperation with the REV7–shieldin complex underpins DNA structure-specific NHEJ. Nature 560, 122–127. doi: 10.1038/s41586-018-0362-1

Gómez-H, L., Felipe-Medina, N., Condezo, Y. B., Garcia-Valiente, R., Ramos, I., Suja, J. A., et al. (2019). The PSMA8 subunit of the spermatoproteasome is essential for proper meiotic exit and mouse fertility. PLoS Genet 15, e1008316. doi: 10.1371/journal.pgen.1008316

Gupta, R., Somyajit, K., Narita, T., Maskey, E., Stanlie, A., Kremer, M., et al. (2018). DNA Repair Network Analysis Reveals Shieldin as a Key Regulator of NHEJ and PARP Inhibitor Sensitivity. Cell 173, 972–988.e23. doi: 10.1016/j.cell.2018.03.050

Han, J., Lee, J.-H., Park, S., Yoon, S., Yoon, A., Hwang, D. B., et al. (2016). A phosphorylation pattern-recognizing antibody specifically reacts to RNA polymerase II bound to exons. Exp Mol Med 48, e271–e271. doi: 10.1038/emm.2016.101

Huang, Y., and Roig, I. (2023). Genetic control of meiosis surveillance mechanisms in mammals. Front Cell Dev Biol 11. doi: 10.3389/fcell.2023.1127440

Humphryes, N., and Hochwagen, A. (2014). A non-sister act: Recombination template choice during meiosis. Exp Cell Res 329, 53–60. doi: 10.1016/j.yexcr.2014.08.024

Hunter, N. (2017). Oocyte Quality Control: Causes, Mechanisms, and Consequences. Cold Spring Harb Symp Quant Biol 82, 235–247. doi: 10.1101/sqb.2017.82.035394

Ishiguro, K., Matsuura, K., Tani, N., Takeda, N., Usuki, S., Yamane, M., et al. (2020). MEIOSIN Directs the Switch from Mitosis to Meiosis in Mammalian Germ Cells. Dev Cell 52, 429–445.e10. doi: 10.1016/j.devcel.2020.01.010

Joyce, E. F., and McKim, K. S. (2010). Chromosome Axis Defects Induce a Checkpoint-Mediated Delay and Interchromosomal Effect on Crossing Over during Drosophila Meiosis. PLoS Genet 6, e1001059. doi: 10.1371/journal.pgen.1001059

Keeney, S., Baudat, F., Angeles, M., Zhou, Z. H., Copeland, N. G., Jenkins, N. A., et al. (1999). A mouse homolog of the Saccharomyces cerevisiae meiotic recombination DNA transesterase Spo11p. Genomics 61, 170–182. Available at: http://www.ncbi.nlm.nih.gov/entrez/query.fcgi?cmd=Retrieve&db=PubMed&dopt=Citation&list_uids=10534402

Kojima, M. L., de Rooij, D. G., and Page, D. C. (2019). Amplification of a broad transcriptional program by a common factor triggers the meiotic cell cycle in mice. Elife 8. doi: 10.7554/eLife.43738

Kumari, N., Kaur, E., Raghavan, S. C., and Sengupta, S. (2025). Regulation of pathway choice in DNA repair after double-strand breaks. Curr Opin Pharmacol 80, 102496. doi: 10.1016/j.coph.2024.102496

Lambing, C., Osman, K., Nuntasoontorn, K., West, A., Higgins, J. D., Copenhaver, G. P., et al. (2015). Arabidopsis PCH2 Mediates Meiotic Chromosome Remodeling and Maturation of Crossovers. PLoS Genet 11, e1005372. doi: 10.1371/journal.pgen.1005372

Mahadevaiah, S. K., Bourc’his, D., de Rooij, D. G., Bestor, T. H., Turner, J. M., and Burgoyne, P. S. (2008). Extensive meiotic asynapsis in mice antagonises meiotic silencing of unsynapsed chromatin and consequently disrupts meiotic sex chromosome inactivation. J Cell Biol 182, 263–276. doi: 10.1083/jcb.200710195; 10.1083/jcb.200710195

Mahadevaiah, S. K., Costa, Y., and Turner, J. M. A. (2009). Using RNA FISH to study gene expression during mammalian meiosis. Methods Mol Biol 558, 433–44. doi: 10.1007/978-1-60761-103-5_25

Mao, Z., Bozzella, M., Seluanov, A., and Gorbunova, V. (2008). Comparison of nonhomologous end joining and homologous recombination in human cells. DNA Repair (Amst) 7, 1765–1771. doi: 10.1016/j.dnarep.2008.06.018

Marcet-Ortega, M., Pacheco, S., Martínez-Marchal, A., Castillo, H., Flores, E., Jasin, M., et al. (2017). p53 and TAp63 participate in the recombination-dependent pachytene arrest in mouse spermatocytes. PLoS Genet 13, e1006845. doi: 10.1371/journal.pgen.1006845

Marco-Sola, S., Sammeth, M., Guigó, R., and Ribeca, P. (2012). The GEM mapper: fast, accurate and versatile alignment by filtration. Nat Methods 9, 1185–1188. doi: 10.1038/nmeth.2221

Martínez-Marchal, A., Huang, Y., Guillot-Ferriols, M. T., Ferrer-Roda, M., Guixé, A., Garcia-Caldés, M., et al. (2020). The DNA damage response is required for oocyte cyst breakdown and follicle formation in mice. PLoS Genet 16, e1009067. doi: 10.1371/journal.pgen.1009067

Mirman, Z., Lottersberger, F., Takai, H., Kibe, T., Gong, Y., Takai, K., et al. (2018). 53BP1–RIF1–shieldin counteracts DSB resection through CST- and Polα-dependent fill-in. Nature 560, 112–116. doi: 10.1038/s41586-018-0324-7

Monesi, V. (1964). Ribonucleic Acid Synthesis during Mitosis and Meiosis in the Mouse Testis. J Cell Biol 22, 521–532.

Montgomery, S. B., Sammeth, M., Gutierrez-Arcelus, M., Lach, R. P., Ingle, C., Nisbett, J., et al. (2010). Transcriptome genetics using second generation sequencing in a Caucasian population. Nature 464, 773–777. doi: 10.1038/nature08903

Nelson, C. R., Hwang, T., Chen, P.-H., and Bhalla, N. (2015). TRIP13PCH-2 promotes Mad2 localization to unattached kinetochores in the spindle checkpoint response. J Cell Biol 211, 503–16. doi: 10.1083/jcb.201505114

Noordermeer, S. M., Adam, S., Setiaputra, D., Barazas, M., Pettitt, S. J., Ling, A. K., et al. (2018). The shieldin complex mediates 53BP1-dependent DNA repair. Nature 560, 117–121. doi: 10.1038/s41586-018-0340-7

Pacheco, S., Marcet-Ortega, M., Lange, J., Jasin, M., Keeney, S., and Roig, I. (2015). The ATM Signaling Cascade Promotes Recombination-Dependent Pachytene Arrest in Mouse Spermatocytes. PLoS Genet 11, e1005017. doi: 10.1371/journal.pgen.1005017

Page, J., de la Fuente, R., Manterola, M., Parra, M. T., Viera, A., Berríos, S., et al. (2012). Inactivation or non-reactivation: what accounts better for the silence of sex chromosomes during mammalian male meiosis? Chromosoma 121, 307–326. doi: 10.1007/s00412-012-0364-y

Paiano, J., Wu, W., Yamada, S., Sciascia, N., Callen, E., Paola Cotrim, A., et al. (2020). ATM and PRDM9 regulate SPO11-bound recombination intermediates during meiosis. Nat Commun 11, 857. doi: 10.1038/s41467-020-14654-w

Pittman, D. L., Cobb, J., Schimenti, K. J., Wilson, L. A., Cooper, D. M., Brignull, E., et al. (1998). Meiotic prophase arrest with failure of chromosome synapsis in mice deficient for Dmc1, a germline-specific RecA homolog. Mol Cell 1, 697–705. Available at: http://www.ncbi.nlm.nih.gov/entrez/query.fcgi?cmd=Retrieve&db=PubMed&dopt=Citation&list_uids=9660953

Rinaldi, V. D., Bloom, J. C., and Schimenti, J. C. (2020). Oocyte Elimination Through DNA Damage Signaling from CHK1/CHK2 to p53 and p63. Genetics 215, 373–378. doi: 10.1534/genetics.120.303182

Rinaldi, V. D., Bolcun-Filas, E., Kogo, H., Kurahashi, H., and Schimenti, J. C. (2017). The DNA Damage Checkpoint Eliminates Mouse Oocytes with Chromosome Synapsis Failure. Mol Cell 67, 1026–1036.e2. doi: 10.1016/j.molcel.2017.07.027

Robinson, M. D., McCarthy, D. J., and Smyth, G. K. (2010). *edgeRL*: a Bioconductor package for differential expression analysis of digital gene expression data. Bioinformatics 26, 139–140. doi: 10.1093/bioinformatics/btp616

Robinson, M. D., and Oshlack, A. (2010). A scaling normalization method for differential expression analysis of RNA-seq data. Genome Biol 11, R25. doi: 10.1186/gb-2010-11-3-r25

Roig, I., Dowdle, J. A. A., Toth, A., de Rooij, D. G. G., Jasin, M., and Keeney, S. (2010). Mouse TRIP13/PCH2 is required for recombination and normal higher-order chromosome structure during meiosis. PLoS Genet 6, e1001062. doi: 10.1371/journal.pgen.1001062

Roig, I., Liebe, B., Egozcue, J., Cabero, Ll., Garcia, M., and Scherthan, H. (2004). Female-specific features of recombinational double-stranded DNA repair in relation to synapsis and telomere dynamics in human oocytes. Chromosoma 113, 22–33. doi: 10.1007/s00412-004-0290-8

Romanienko, P. J., and Camerini-Otero, R. D. (2000). The mouse Spo11 gene is required for meiotic chromosome synapsis. Mol Cell 6, 975–987.

Royo, H., Polikiewicz, G., Mahadevaiah, S. K., Prosser, H., Mitchell, M., Bradley, A., et al. (2010). Evidence that meiotic sex chromosome inactivation is essential for male fertility. Curr Biol 20, 2117–2123. doi: 10.1016/j.cub.2010.11.010

San-Segundo, P. A., and Roeder, G. S. (1999). Pch2 links chromatin silencing to meiotic checkpoint control. Cell 97, 313–324. Available at: http://www.ncbi.nlm.nih.gov/entrez/query.fcgi?cmd=Retrieve&db=PubMed&dopt=Citation&list_uids=10319812

Sarangi, P., Clairmont, C. S., Galli, L. D., Moreau, L. A., and D’Andrea, A. D. (2020). p31 ^comet^ promotes homologous recombination by inactivating REV7 through the TRIP13 ATPase. Proceedings of the National Academy of Sciences 117, 26795–26803. doi: 10.1073/pnas.2008830117

Shin, Y.-H., Choi, Y., Erdin, S. U., Yatsenko, S. A., Kloc, M., Yang, F., et al. (2010). Hormad1 mutation disrupts synaptonemal complex formation, recombination, and chromosome segregation in mammalian meiosis. PLoS Genet 6, e1001190. doi: 10.1371/journal.pgen.1001190

Takai, H., Naka, K., Okada, Y., Watanabe, M., Harada, N., Saito, S., et al. (2002). Chk2-deficient mice exhibit radioresistance and defective p53-mediated transcription. EMBO J 21, 5195–5205.

Turner, J. M., Aprelikova, O., Xu, X., Wang, R., Kim, S., Chandramouli, G. v, et al. (2004). BRCA1, histone H2AX phosphorylation, and male meiotic sex chromosome inactivation. Curr Biol 14, 2135–2142. Available at: http://www.ncbi.nlm.nih.gov/entrez/query.fcgi?cmd=Retrieve&db=PubMed&dopt=Citation&list_uids=15589157

Wojtasz, L., Cloutier, J. M., Baumann, M., Daniel, K., Varga, J., Fu, J., et al. (2012). Meiotic DNA double-strand breaks and chromosome asynapsis in mice are monitored by distinct HORMAD2-independent and -dependent mechanisms. Genes Dev 26, 958–973. doi: 10.1101/gad.187559.112; 10.1101/gad.187559.112

Wojtasz, L., Daniel, K., Roig, I., Bolcun-Filas, E., Xu, H., Boonsanay, V., et al. (2009). Mouse HORMAD1 and HORMAD2, two conserved meiotic chromosomal proteins, are depleted from synapsed chromosome axes with the help of TRIP13 AAA-ATPase. PLoS Genet 5, e1000702. doi: 10.1371/journal.pgen.1000702

Yamada, S., Hinch, A. G., Kamido, H., Zhang, Y., Edelmann, W., and Keeney, S. (2020). Molecular structures and mechanisms of DNA break processing in mouse meiosis. Genes Dev 34, 806–818. doi: 10.1101/GAD.336032.119

Yoshida, K., Kondoh, G., Matsuda, Y., Habu, T., Nishimune, Y., and Morita, T. (1998). The mouse RecA-like gene Dmc1 is required for homologous chromosome synapsis during meiosis. Mol Cell 1, 707–718. Available at: http://www.ncbi.nlm.nih.gov/entrez/query.fcgi?cmd=Retrieve&db=PubMed&dopt=Citation&list_uids=9660954

