## Supplementary tables for "TRIP13 fosters both transcriptional silencing and DSB repair during meiosis"

**Table S1.** Correlation of the number of genes per chromosome and the number of upregulated and downregulated genes in *Trip13^mod/mod^* testis.

| Chromosome | Number genes | Percentage genes | Number upregulated genes | Percentage upregulated genes | Number downregulated genes | Percentage downregulated genes |
| --- | --- | --- | --- | --- | --- | --- |
| 1 | 1215 | 5.4% | 3 | 1.8% | 23 | 7.0% |
| 2 | 1851 | 8.3% | 3 | 1.8%* | 23 | 7.0% |
| 3 | 1009 | 4.5% | 4 | 2.5% | 14 | 4.3% |
| 4 | 1313 | 5.9% | 5 | 3.1% | 16 | 4.9% |
| *5* | 1263 | 5.6% | 2 | 1.2%* | 26 | 8.0% |
| *6* | 1265 | 5.6% | 1 | 0.6%* | 19 | 5.8% |
| *7* | 2038 | 9.1% | 3 | 1.8%* | 20 | 6.1% |
| *8* | 1035 | 4.6% | 3 | 1.8% | 22 | 6.7% |
| *9* | 1221 | 5.4% | 4 | 2.5% | 15 | 4.6% |
| *10* | 1013 | 4.5% | 4 | 2.5% | 18 | 5.5% |
| *11* | 1632 | 7.3% | 3 | 1.8%* | 25 | 7.6% |
| *12* | 785 | 3.5% | 3 | 1.8% | 8 | 2.4% |
| *13* | 849 | 3.8% | 7 | 4.3% | 16 | 4.9% |
| *14* | 1068 | 4.8% | 1 | 0.6%* | 8 | 2.4% |
| *15* | 785 | 3.5% | 0 | 0.0%* | 20 | 6.1%* |
| *16* | 681 | 3.0% | 0 | 0.0%* | 8 | 2.4% |
| *17* | 1064 | 4.7% | 1 | 0.6%* | 27 | 8.3%* |
| *18* | 515 | 2.3% | 2 | 1.2% | 9 | 2.8% |
| *19* | 722 | 3.2% | 1 | 0.6%* | 10 | 3.1% |
| *X* | 933 | 4.2% | 106 | 65.0%* | 0 | 0.0%* |
| *Y* | 172 | 0.8% | 7 | 4.3%* | 0 | 0.0% |

* Denotes p<0.05; Chi-square test with Yates correction

**Table S2.** Oocyte number and follicle type in 1-month-old mouse ovaries. The numbers express the average ± SEM. N indicates the number of ovaries counted.

| Genotype | N | Total number of oocytes | Primordial  Follicles | Primary Follicles | Secondary  Follicles | Antral Follicles | Abnormal  Follicles |
| --- | --- | --- | --- | --- | --- | --- | --- |
| Wild type* | 7 | 874.9 ± 64.2 | 629.3 ± 68.0 | 147.1 ± 17.2 | 71.4 ± 3.8 | 20.43 ± 3.6 | 6.6 ± 0.6 |
| *Spo11^-/-^* | 8 | 67.0 ± 12.1 | 2.8 ± 0.9 | 2.0 ± 7.4 | 32.9 ± 7.4 | 24.9 ± 3.9 | 4.5 ± 1.2 |
| *Spo11^-/-^ Trip13^mod/mod^* | 8 | 83.0 ± 13.0 | 2.0 ± 1.2 | 3.9 ± 1.5 | 37.8 ± 7.8 | 30.6 ± 2.8 | 8.8 ± 2.3 |
| *Dmc1^-/-^ Chk2^-/-^* | 8 | 366.6 ± 16.3 | 108.4 ± 11.8 | 59.9 ± 7.2 | 127.0 ± 4.6 | 42.1 ± 2.9 | 28.9 ± 4.2 |
| *Dmc1^-/-^ Chk2^-/-^ Trip13^mod/mod^* | 8 | 648.9 ± 110.8 | 366.5 ± 84.4 | 109.9 ± 11.1 | 123.6 ± 15.4 | 32.9 ± 4.8 | 16.0 ± 1.8 |

*Data from Martínez-Marchal et al., 2020.
